# CD39 Identifies Tumor-Reactive CD8 T cells in Patients With Lung Cancer

**DOI:** 10.1101/2022.01.24.477554

**Authors:** Andrew Chow, Fathema Z. Uddin, Levi Mangarin, Hira Rizvi, Anton Dobrin, Sam Tischfield, Alvaro Quintanal-Villalonga, Joseph M. Chan, Nisargbhai Shah, Viola Allaj, Parvathy Manoj, Marissa Mattar, Maximiliano Meneses, Michael Liu, Rebecca Landau, Mariana Ward, Amanda Kulick, Charlene Kwong, Matthew Wierzbicki, Jessica Yavner, Shweta S. Chavan, Abigail Farillas, Aliya Holland, Harsha Sridhar, Metamia Ciampricotti, Daniel Hirschhorn, Allison L Richards, Mark T.A. Donoghue, Glenn Heller, Christopher A. Klebanoff, Matthew D. Hellmann, Elisa de Stanchina, Triparna Sen, Jedd D. Wolchok, Taha Merghoub, Charles M. Rudin

## Abstract

The repertoire of tumor-infiltrating lymphocytes (TILs) can be vast, and many of these TILs are not endowed with tumor reactivity. While a number of reports have shown that tumor-reactive TILs express CD39, few reports have demonstrated that conversely, CD39 can be leveraged to serve as a proxy of tumor-reactive CD8 T cells. Using single-cell CITE/RNA/TCRseq, we show that CD39^+^ CD8 T cells in human lung cancers demonstrate transcriptional and proteomic features of exhaustion, tumor reactivity, and clonal expansion. Moreover, TCR cloning revealed that CD39 enriched for tumor-reactive CD8 T cell clones. Flow cytometry of 440 lung cancer specimens revealed that CD39 level on CD8 T cells is only weakly correlated with tumoral features that currently guide lung cancer therapy, such as histology, driver mutation, PD-L1 and tumor mutation burden. PD-1 axis blockade, but not cytotoxic chemotherapy, increased intratumoral CD39^+^ CD8 T cells. CD39 correlated with PD-1 expression on CD8 T cells and high pre-treatment/early-on-treatment levels were associated with improved clinical outcomes, but not immune-related adverse events, from immune checkpoint blockade therapy. This comprehensive profiling of the clinical, pathological and molecular features highlights the utility of CD39 as a proxy for tumor-reactive CD8 T cells in human lung cancer.

## INTRODUCTION

Lung cancer is the leading cause of cancer death in the world. Immune checkpoint blockade (ICB) utilizing antibodies that block programmed cell death protein 1 (PD-1), programmed death-ligand 1 (PD-L1), and cytotoxic T-lymphocyte-associated protein 4 (CTLA-4) has ushered in a new era of hope, with long-term durable disease control in a subset of patients with lung cancer (*1, 2*). However, most patients do not respond to ICB therapy and many of those who initially respond eventually develop recurrent and progressive disease. While PD-L1 on tumor cells and tumor mutation burden (TMB) have been validated as predictors of benefit from ICB in lung cancer (*3, 4*), there is an opportunity to further refine these biomarkers by incorporating features of CD8 T cells, which are critical mediators of ICB efficacy.

Despite early suggestions that measurement of bulk CD8 tumor infiltrating lymphocytes (TILs) would predict benefit from ICB (5), multiple recent studies across different diagnostic modalities have demonstrated that total CD8 T cell content alone is not a reliable predictor of clinical benefit from ICB (6-10). Part of the reason for this lack of predictive benefit is that a large proportion of CD8 T cells in the tumor microenvironment are passive bystanders that lack tumor reactivity (*11, 12*). Recent elegant work has highlighted that CD8 T cells that express CD39 are enriched for features of a clonal, proliferative lymphocyte population that expresses high levels of activation/exhaustion (e.g. PD-1) and cytotoxicity (e.g. granzyme B) markers (*2, 12-15*). Moreover, CD39 was highly expressed on empirically defined neoantigen-and tumor-associated antigen-reactive T cells in lung cancer and melanoma (*16-19*). The largest profiling of CD39 expression on CD8 T cells to date in a single disease was performed in 68 patients with head and neck squamous cell cancer; this paper contained an additional 42 samples across a mixture of seven disease types (13). Herein, we describe the tumoral features that are associated with CD39 levels on CD8 T cells from 440 clinical samples of human lung cancer, validate the utility of CD39 as a surrogate of tumor-reactive CD8 T cells, and report that baseline CD39 on CD8 T cells predicts enhanced efficacy of ICB in patients with lung cancer.

## METHODS

### Human biospecimens

Fresh primary tumors, metastatic lesions and pleural/peritoneal/pericardial effusions were obtained from August 2018 to September 2021 with permission from the MSKCC IRB. Informed consent was collected from all patients enrolled in this study. Clinical samples were annotated with tumor histology and driver mutation. Adenocarcinoma, adenosquamous, and NSCLC NOS tumors were annotated by their molecular driver mutations, if known. The category ‘Unknown Driver’ refers to adenocarcinoma, adenosquamous, and NSCLC NOS histology tumors for which a driver mutation (defined as ‘known to be oncogenic’ by OncoKB (*20*)) was not identified; notably, this category does not include squamous or small-cell histology biospecimens. With the exception of a single case in which a tumor sample with squamous histology harbored a *MET* exon 14 mutation and two transformed small-cell lung cancer tumors with *EGFR* mutations, squamous and small-cell lung cancers were annotated by their histology.

### Flow cytometry

Cells were incubated with TruFCX (Biolegend) to block nonspecific binding, and then stained (15 min, 4 °C) with appropriate dilutions of various combinations of fluorochrome-conjugated anti-mouse antibodies. The stained cells were acquired on a LSRII Flow Cytometer or Aria 7 cell sorter and the data were processed using FlowJo software (Treestar). Doublets and dead cells were excluded based on forward scatter (FSC) and side scatter (SSC) and 4′,6-diamidino-2-phenylindole staining (DAPI, 1 µg/ml; ThermoFisher). All depicted flow cytometry plots were pre-gated on non-debris (by FSC and SSC), viable (DAPI^-^) single CD45^+^ CD3^+^ cells, unless otherwise indicated in the Figure legend. Gating for %CD39 was determined by gating of fluorescence minus one (FMO). PD-1 mean fluorescence intensity was measured for a subset of samples.

### Single-cell transcriptome sequencing

Sorted or dissociated tumor cells were stained with Trypan blue and Countess II Automated Cell Counter (ThermoFisher) was used to assess both cell number and viability. Following QC, the single-cell suspension was loaded onto Chromium Chip A or Next GEM Chip K (10X Genomics PN 230027/1000286) and GEM generation, cDNA synthesis, cDNA amplification, and library preparation of 700-3,300 cells proceeded using the Chromium Single Cell 5’ Reagent Kit or Next GEM Single Cell 5’ Kit v2 (10X Genomics PN 1000006/1000263) according to the manufacturer’s protocol. cDNA amplification included 14-16 cycles and 5.8 ng-20 ng of the material was used to prepare sequencing libraries with 16 cycles of PCR. Indexed libraries were pooled equimolar and sequenced on a NovaSeq 6000 in a PE26/91 or PE28/91 run using the NovaSeq 6000 SP or S1 Reagent Kit (100 cycles) (Illumina). An average of 108 million reads were generated per sample.

### Single-cell V(D)J analysis from RNA

An aliquot of complementary DNA (cDNA) generated using the methods described above was used to enrich for V(D)J regions using the Chromium Single Cell V(D)J Enrichment Kit Human T Cell (10X Genomics PN 1000005) according to the manufacturer’s protocol with 10 cycles of PCR during enrichment and 9 cycles during library preparation. Indexed libraries were pooled equimolar and sequenced on a NovaSeq 6000 in a PE150 or PE26/91 run using the NovaSeq 6000 SP or S4 Reagent Kit (100 or 300 cycles) (Illumina). An average of 24 million paired reads was generated per sample.

### Cell surface protein feature barcode analysis

Amplification products generated using the methods described above included both cDNA and feature barcodes tagged with cell barcodes and unique molecular identifiers. Smaller feature barcode fragments were separated from longer amplified cDNA using a 0.6X cleanup with aMPure XP beads (Beckman Coulter catalog # A63882). Libraries were constructed using the Chromium Single Cell 5’ Feature Barcode Library Kit (10X Genomics PN 1000080) according to the manufacturer’s protocol with 9 cycles of PCR. Indexed libraries were pooled equimolar and sequenced on a NovaSeq 6000 in a PE26/91 or PE28/91 run using the NovaSeq 6000 SP or S2 Reagent Kit (100 cycles) (Illumina). An average of 60 million paired reads was generated per sample.

### Single cell CITE/RNA/TCR analysis

Single-cell sequencing data were aligned to the Genome Reference Consortium Human Build 38 (GRCh38) using Cell Ranger (v3.1.0; 10X Genomics) in order to obtain T cell clonotypes, feature barcoding, CITEseq antibody detection and gene expression profiles associated with individual single cells. Each data type was matched to create a UMI matrix and cells were filtered out based on three metrics: (1) cells with fewer than 200 detectable genes; (2) cells with more than 3000 detectable genes; (3) cells that had fewer than 5% percentage of counts related to mitochondrial genes. Data normalization, Principal Component Analysis and subsequent Uniform Manifold Approximation and Projections (UMAP) were performed on the dataset using the R package Seurat v.3.1.1 (https://github.com/satijalab/seurat). The differential expression comparisons were generated using the DESeq2 package with selected genes (FDR < 0.05). After filtering, we created subclusters of cells using the 26 Louvain algorithm. Raw counts were normalized by library size per cell. CD39^neg^ was defined as a normalized value of 0 for adt_CD39. CD39^int^ was defined as the adt_CD39 level between 0 and 1.0 (non-inclusive) since 1.0 was the mean of all non-zero values for adt_CD39 among all the cells in the CD8 cluster. CD39hi was defined as adt_CD39 level greater than or equal to 1.0. Signature scores for exhaustion and T cell reactivity were calculated by *AddModuleScore* in Seurat. Clonal proportion was calculated as the fractional representation of all CD8 clones by clonotypes that were categorized by mean adt_CD39 expression with the same cutoffs for CD39^low^, CD39^int^, and CD39^hi^ as above.

### Multivariate analyses

We used the R package caret (*21*) to implement the glmnet (*22*) algorithm and evaluate the performance of a lasso regression model using 10-fold cross-validation. The data was center and scaled during preprocessing and lambda was chosen based on the minimal RMSE value. Additionally a linear model was evaluated using the R function “lm”. In both models, the response variable, CD39 expression, was log base 2 transformed as well as the predictor variables TMB and total neoantigens per sample.

### Clinical outcomes analyses

RFS assessment was performed on 188 biospecimens obtained from stage I-IIIA lung cancer who did not receive neoadjuvant ICB and were not lost to follow up after resection. For the PFS assessment in stage IV patients not treated with ICB, only patients that received at least two cycles of platinum-based chemotherapy were included in the analysis (n=26). For the PFS assessment in stage IV patient treated with ICB, only patients that received at least two cycles of ICB without chemotherapy were included in the analysis (n=22). The cutoffs for %CD8 and %CD39 of 15.4 and 23.3 were selected from the means of the 440-sample cohort. The PD-1 cutoff was selected to divide the cohort above and below the median for the stage IV cohort. Response criteria were annotated per RECIST v1.1.

### Statistical analysis

Data are expressed and statistical analyses were performed as described in the Figure legend for each analysis. Statistical significance was determined by two-way ANOVA with Tukey’s multiple comparison test, student’s t-test or Mantel-Cox log-rank test using Prism 7 software as indicated.

### Materials availability

This study did not generate new unique reagents.

### Data availability

All data generated and supporting the findings of this study are available within the paper. The single-cell CITE/TCR/RNAseq will be deposited in the Gene Expression Omnibus (GEO) database. The TCRseq data will be available through ImmuneACCESS on the Adaptive Biotechnologies website.

## RESULTS

### Single-cell cellular indexing of transcriptomes and epitopes (CITE)/RNA/T cell receptor (TCR)seq reveals that CD39^high^ CD8 T cells are enriched for features of exhaustion, tumor reactivity, and clonal expansion

To characterize CD8 T cells expressing CD39, we sorted CD3^+^ T cells from four non-small cell lung cancer (NSCLC) biospecimens and performed droplet-based single-cell CITE/RNA/TCRseq (*23*). The four NSCLC samples comprised a range of histologies (e.g. adenocarcinoma and squamous), driver mutations (*KRAS* and *EGFR*), stages (e.g. early and metastatic), and anatomic sites (e.g. lung, pleural fluid, and brain) (**Table S1**). None of these patients had received prior ICB at the time of sample collection. After coarse clustering, we visualized separated clusters of regulatory T cells, CD4 T cells, CD8 T cells, and an additional T cell NOS (not otherwise specified) cluster (**Fig. 1A, Table S2**). We focused our subsequent analyses on CD8 T cells since this subset has the greatest known contribution to anti-tumor immunity. The 896 single CD8 T cells that passed quality control (QC) were divided by CD39 protein expression (assessed by oligo-tagged anti-CD39 antibody) into high (hi), intermediate (int), and negative (neg) groups. Transcriptional dropout is a well-known limitation of single cell RNAseq, and oligonucleotide-tagged antibodies to surface molecules (e.g. CITE-seq) represents a strategy to overcome this hurdle (*24*). Concordantly, CITEseq detected protein expression of CD39 and other activation markers in many cells in which there was absent transcriptional expression (**Fig S1A**). Genes that were differentially expressed in CD39^hi^ CD8 T cells included the exhaustion marker *LAYN*, tissue residence marker *ITGAE* (CD103), and the activation markers *CXCL13, GNLY, HLADRA*, and *VCAM1* (**Table S3**). CD39^hi^ CD8 T cells had the highest transcriptional levels of *ENTPD1* (CD39), *PDCD1* (PD-1), *ITGAE, CXCL13, TNFRSF4* (OX-40), *HAVCR2* (TIM-3), and *LAG3*, which are features of tumor-reactive CD8 T cells (**Fig. S1B**) (*25*). Protein level assessment by CITE-seq revealed that CD39^hi^ CD8 T cells across the four samples consistently expressed the highest protein levels of PD-1, CD103, OX40, 4-1BB, and LAG-3 (**Fig. 1B**). Concordantly, CD39^hi^ CD8 T cells expressed the highest transcriptomic signature score for T cell exhaustion and tumor reactivity (**Fig. 1C-D, Table S4**).

**Figure 1.**
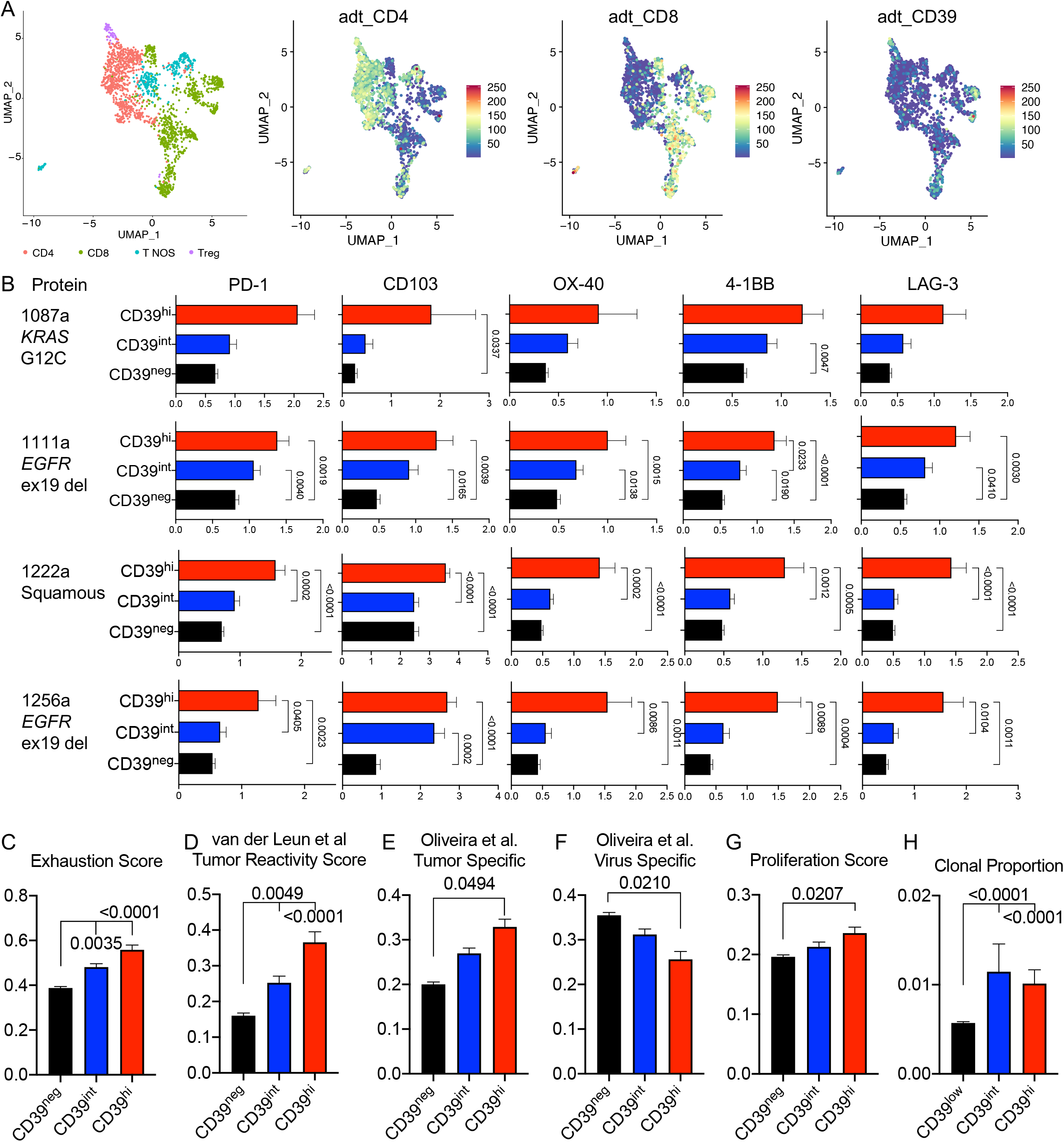
Single-cell CITE/RNA/TCR-sequencing reveals that CD39^hi^ CD8 T cells are enriched for features of exhaustion, tumor-reactivity, and clonal proliferation in human lung cancer. A) UMAP of sorted CD3^+^ T cells from four patients with lung cancer (see Table S1 for patient characteristics). Clusters are annotated on left panel. Surface levels of CD4, CD8, and CD39 as assessed by CITEseq antibody-derived tags (adt) are depicted in right three panels. B) Levels of various proteins (each column) across the four samples (each row) as determined by CITEseq adt levels. C-G) Scaled scores for exhaustion, tumor reactivity, tumor specific, virus specific, and proliferation gene signatures (see Table S4 for full gene list). H) Clonal proportion among CD8 T cells of clonotypes that were categorized by mean CD39 expression. Statistical significance was determined with two-way ANOVA with Tukey’s multiple comparisons test and p value is indicated if < 0.05.

Two recent reports demonstrated that empirically validated tumor-reactive CD8 T cells express CD39 (*16, 17*). In these studies, the authors assessed for CD8 tumor reactivity to tumor-associated or viral-associated antigens and generated gene expression signatures of the reactive CD8 T cells. Thus, we next evaluated whether the gene expression profiles of CD39^hi^ CD8 T in our dataset overlapped with the CD8 T cells in these two recent datasets. CD39^hi^ CD8 T cells in our dataset indeed expressed the highest ‘tumor specific’ and “MANA-TIL (mutation-associated neoantigen-tumor infiltrating lymphocyte)’ signature scores and lowest ‘virus specific’ and ‘influenza’ signatures (**Fig. 1E-F, S1C-D**).

Since CD39^hi^ CD8 T cells were enriched for features of tumor reactivity, we hypothesized that these cells would display greater clonal expansion. Indeed CD39^hi^ CD8 T cells expressed the highest proliferation score and both CD39^int^ and CD39^hi^ CD8 T cells comprised a higher clonal proportion among all CD8 T cells (**Fig 1G-H**). Thus, our single cell CITE/RNA/TCR sequencing demonstrates that CD39^hi^ CD8 T cells express features of exhaustion, tumor reactivity, and clonal expansion.

### CD39 expression on CD8 T cells is a non-redundant biomarker

We compared CD39, Tim-3, 4-1BB, and PD-1 expression by flow cytometry to compare their relative staining resolution, as defined by the separation of the positive and negative population. FACS-based detection of CD39 consistently yielded higher resolution compared to the other three markers (**Fig. 2A**). The enhanced resolution of CD39 compared to PD-1 is consistent with our prior report that CD39 has more durable expression in CD8 T cells (*26*). Due to the staining resolution of the CD39 marker and its association with tumor reactivity, we sought to utilize CD39 expression on CD8 T cells to estimate the frequency of tumor-reactive CD8 T cells in lung cancer clinical samples. From August 2018 to September 2021, we evaluated 440 fresh lung cancer clinical specimens by flow cytometry for CD39 expression on CD8 T cells. These biospecimens ranged across stages (I-IV) and lung cancer subtypes (lung adenocarcinoma, squamous cell cancer, and small cell lung cancer) (**Table S5**). The mean CD8 level was 15.4% (of CD45^+^) and mean CD39 level was 23.3% (of CD8 T cells), and these were utilized as cutoffs throughout the rest of the study. On univariate analysis, CD39 expression on CD8 T cells had a weak correlation with total CD8 T cells, smoking history, TMB, and PD-L1 (**Fig. S2A-D**). Since TMB ≥ 10 mutations/megabase and PD-L1 ≥ 50% represent subgroups with favorable clinical outcomes from ICB therapy (*3, 4*), we assessed %CD39 on these subpopulations. When divided into quartiles by PD-L1 and TMB expression, tumor material from the two TMB ≥ 10 mutations/megabase subgroups (both PD-L1 < 50% and ≥ 50%) had nearly two-fold higher %CD39 among CD8 T cells, relative to the TMB < 10 and PD-L1 < 50% subgroup (**Fig. S2E**). In contrast, the proportion of CD8 T cells (**Fig. S2F**) showed less variation. There was largely no association of CD39 expression on CD8 T cells with lung cancer stage with the exception of a reduction in stage IVA tumors (**Fig. S2G**). This is concordant with reduced CD39 expression on CD8 T cells in pleural fluid and pleural metastases biospecimens relative to the lung biospecimens (**Fig. S2H**). Notably, we previously reported that the pleural and peritoneal cavities are immunosuppressed microenvironments due to the presence of Tim-4^+^ cavity-resident macrophages (*26*), and this may contribute to the low %CD39 in these anatomic compartments.

**Figure 2.**
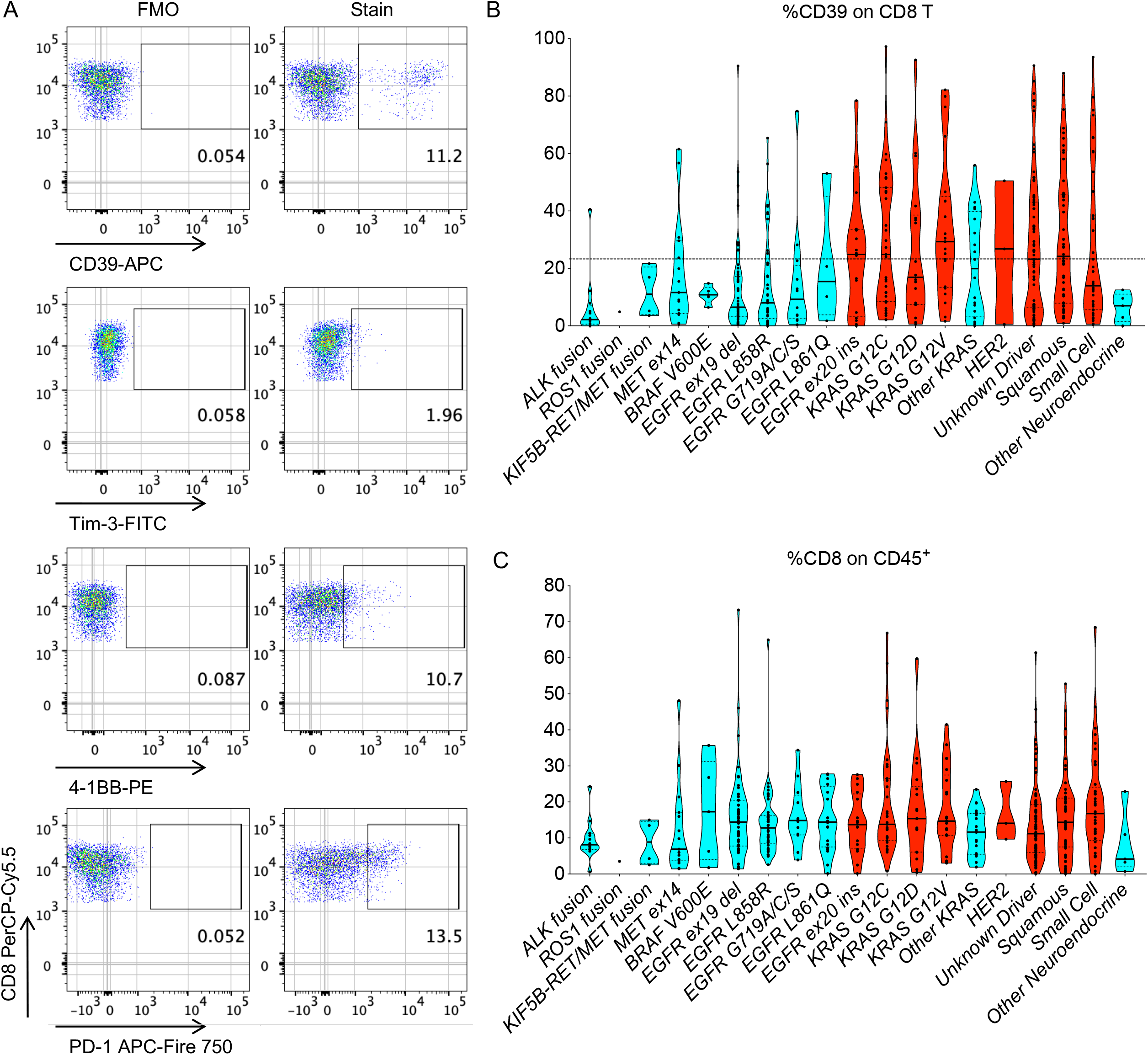
CD39 expression among CD8 TILs varies with lung cancer subtype. A) Representative flow cytometry staining of CD39, Tim-3, 4-1BB, and PD-1 on DAPI^-^ CD45^+^ CD3^+^ CD8^+^ T cells from MSK 1105b. Left plot represents fluorescence minus one (FMO) and right plot represents the CD8 T cells stained with the indicated antibody. B-C) Violin plots of %CD39^+^ (among CD8^+^ T cells) and %CD8 (among CD45^+^) for various histological subtypes/driver mutation categories among the 440-sample cohort. Dashed line indicates 23.3% mean CD39 level for entire 440-sample cohort. Red bars indicate histological subtypes/driver mutations with above average %CD39 values.

CD39 levels on CD8 T cells showed two patterns of expression among the lung cancer subtypes. *ALK* fusion, *ROS1* fusion, *MET* exon 14 fusion, *BRAF* V600E, *RET* fusion and *EGFR* mutant (except exon 20 insertion) lung adenocarcinomas, which are not associated with tobacco use, tended to have below average CD39 expression on CD8 T cells (**Fig. 2B, Table S6**). The reduced CD39 expression on *EGFR* mutant lung cancer tumors is in line with a prior report (*12*). In contrast, CD39 expression on lung adenocarcinomas with *KRAS* G12C, G12D, G12V or *ERBB2* (HER2) mutations were above average. ‘Other *KRAS*‘ mutations, including G12A, G12R, G12S, G13C, G13D, KRAS Q61H, and Q61R, which are known to be oncogenic per OncoKB (*20*), were associated with a lower-than-mean CD39 level on CD8 T cells. The proportion of CD39^+^ CD8 T cells in squamous and small cell lung cancers averaged 30.3% and 27.3%, respectively. In contrast, only 6.4% of CD8 T cells in neuroendocrine carcinoid (other neuroendocrine) tumors expressed CD39. These findings are consistent with *KRAS*, squamous, and small-cell lung cancers being strongly associated with tobacco use, which is correlated with a higher number of tumor mutations and neoantigens to which CD8 T cells can react. Nonetheless, even among the lung cancer subtypes with higher %CD39, there was a wide range of CD39 levels (**Fig. 2B**), suggesting that even for a given driver mutation, there is substantial heterogeneity in tumor-reactive CD8 immunosurveillance. Total CD8 T cell infiltration, TMB, and PD-L1 were overall not associated with consistent differences across lung cancer histological subtypes (**Fig. 2C, S2I-J**).

Since our data was retrospective and derived from a heterogeneous cohort of patients, we performed multivariate analyses to determine the clinical and molecular features that best correlate with CD39 expression on CD8 T cells. We included tissue site, stage, histology, driver mutation, TMB, smoking history, and PD-L1 as potential covariates. We also included human leukocyte antigen (HLA) heterozygosity (*27, 28*) in the model and also the number of predicted neoantigens and strong binding neoantigens from next-generation sequencing of tumor biopsies by MSK-IMPACT (*29*). A linear model predicted the variance in %CD39 on CD8 T cells at an adjusted R-squared of 0.24, suggesting that the features in the model poorly predict the variance observed in CD39 levels (**Table S7**). In the model, only TMB, PD-L1, pericardial metastases, and prior chemotherapy passed significance thresholds of *P* < 0.05. Since some of the features in the dataset had few observations and to avoid overfitting, we also applied a lasso regression with 10-fold cross validation. The lambda was selected based on the root-mean-square error (RMSE) and the model achieved an R-squared of 0.17 with a RMSE of 1.5 at lambda = 0.1 (**Table S8**). Notably, at higher lambdas, the algorithm removed all predictors suggesting that there was no linear combination of any regressed parameters that predicted CD39 levels well. Overall, these results indicate that CD39 expression on CD8 T cells is a feature of the tumor that is non-redundant to the tumoral parameters that currently guide therapy in lung cancer (e.g. histology, driver alterations PD-L1, and TMB).

### CD39 is upregulated on CD8 T cells in the tumor microenvironment

We next sought to understand the patterns of tissue-specific expression for CD39 and thus performed CD39 staining on CD8 T cells from matched tumor and peripheral blood. We analyzed three paired samples from peripheral blood and matched tumor lesions with varied anatomic sites (e.g. the brain, lymph node, and lung). In all three cases, high levels of CD39 were only observed in the tumor tissue and not in the peripheral blood (**Fig. S3A**). This suggests that CD8 T cell interactions with cancer cells drives the expression of CD39 and is in line with our recent report that CD39 was expressed at higher levels in regions of a resected tumor with viable cancer cells compared to regions without viable tumor, normal adjacent regions, and lymph nodes (*30*). Since there was a small CD39^+^ population of intermediate intensity in the peripheral blood, we evaluated whether this population gave rise to the CD39^+^ CD8 T cells in the tumor tissue. To address this question, we sorted CD39^-^ and CD39^+^ CD8 T cells from the peripheral blood and CD39^-^ or CD39^+^ from the tumor tissue. We then performed bulk TCR sequencing analysis and assessed for clonal overlap. As assessed by the Morisita Index, there was a minimal level of clonal overlap between CD39^+^ CD8 T cells from the peripheral blood and CD39^+^ CD8 T cells from tumor tissue, suggesting that circulating CD39^+^ CD8 T cells may not be the dominant precursors for the CD39^+^ CD8 T cells in the tumor microenvironment (**Fig. S3A-C**).

We have previously shown that CD39 surface expression manifests more durable expression after TCR stimulation compared to other markers associated with activation, such as phosphatidylserine, PD-1, and CD44 (*26*). Due the greater durability of CD39 even after removal of TCR stimulation, we next assessed the robustness of CD39 levels across spatially and temporally distinct lesions. We analyzed 12 paired samples in which two anatomically distinct lesions were simultaneously assessed (e.g. two resected lung lesions, lymph node and primary lung lesions, or pleural metastasis and fluid). We noted reasonable concordance across spatially distinct lesions (**Fig. S3D**), as exemplified by the low CD39 level on CD8 T cells from the paired pleural metastasis and fluid from MSK 1266 (**Fig. S3E**). There was one notable case of divergent CD39 levels. MSK 1372a and b were simultaneous resections of two right-sided separate primary lung cancer lesions with marked differences in CD39 expression (59.0% vs. 92.5%; **Fig. S3F**). Consistent with being separate primary lesions, these two tumors were comprised of two different *KRAS* driver mutations with differential levels of TMB, which may underly the divergence of CD39 expression. We also followed CD39 levels on CD8 T cells from the same anatomic site multiple times (e.g. recurrent pleural effusions, initial lung biopsy followed by resection, and serial resections for recurrent disease in the brain). The CD39 levels over time across nine patients were largely durable, including for three brain lesions that were resected over the course of nine months (MSK 1265; **Fig. S3G-H**). These findings highlight that CD39 is upregulated on CD8 T cells in proximity to the tumor and that CD39 levels on CD8 T cells are relatively preserved across spatially and temporally distinct lesions in the same patient.

### PD-1 axis blockade increases the frequency of CD39^+^ CD8 T cells

Since our data and prior reports (*12, 16*) demonstrate that tumor antigen-reactive CD8 T cells express CD39 in lung cancer, we sought to leverage CD39 as a proxy for CD8 anti-tumor immunosurveillance. It has been proposed that chemotherapy can result in ‘immunogenic cell death’ that can prime an anti-tumor CD8 T cell response (*31*). We reasoned that CD39 levels would be increased in patients after cytotoxic chemotherapy if such immune priming occurred. Analogously, ICB therapy has been described to expand the pool of antigen-specific CD8 T cells that infiltrate the tumor (*32*), which we reasoned might be reflected in increased CD39^+^ CD8 T cells in the tumor. Thus, we sought to evaluate in our dataset whether we found increases of CD39^+^ CD8 T cells with either cytotoxic chemotherapy or ICB therapy.

We first investigated the 218 patients in our cohort with stage IV lung cancer who were treated with or without chemotherapy and/or ICB in the prior three or six months. Both total CD8 infiltration and CD39 levels on CD8 T cells were unchanged in patients receiving cytotoxic chemotherapy in the preceding three or six months (**Fig. 3A-D**). Total CD8 infiltration was also unchanged in patients receiving preceding ICB therapy (**Fig. 3A,C**). In contrast, CD8 T cells from patients that received ICB in the prior three or six months expressed higher levels of CD39 (**Fig. 3B,D**). In patients that received both cytotoxic chemotherapy and ICB therapy in prior three or six months, the level of CD39 on CD8 T cells was unchanged compared to patients that had not received chemotherapy or ICB in the prior 3 or 6 months, which is consistent with chemotherapy and ICB having opposing effects on CD39 expression (**Fig. 3B,D**).

**Figure 3.**
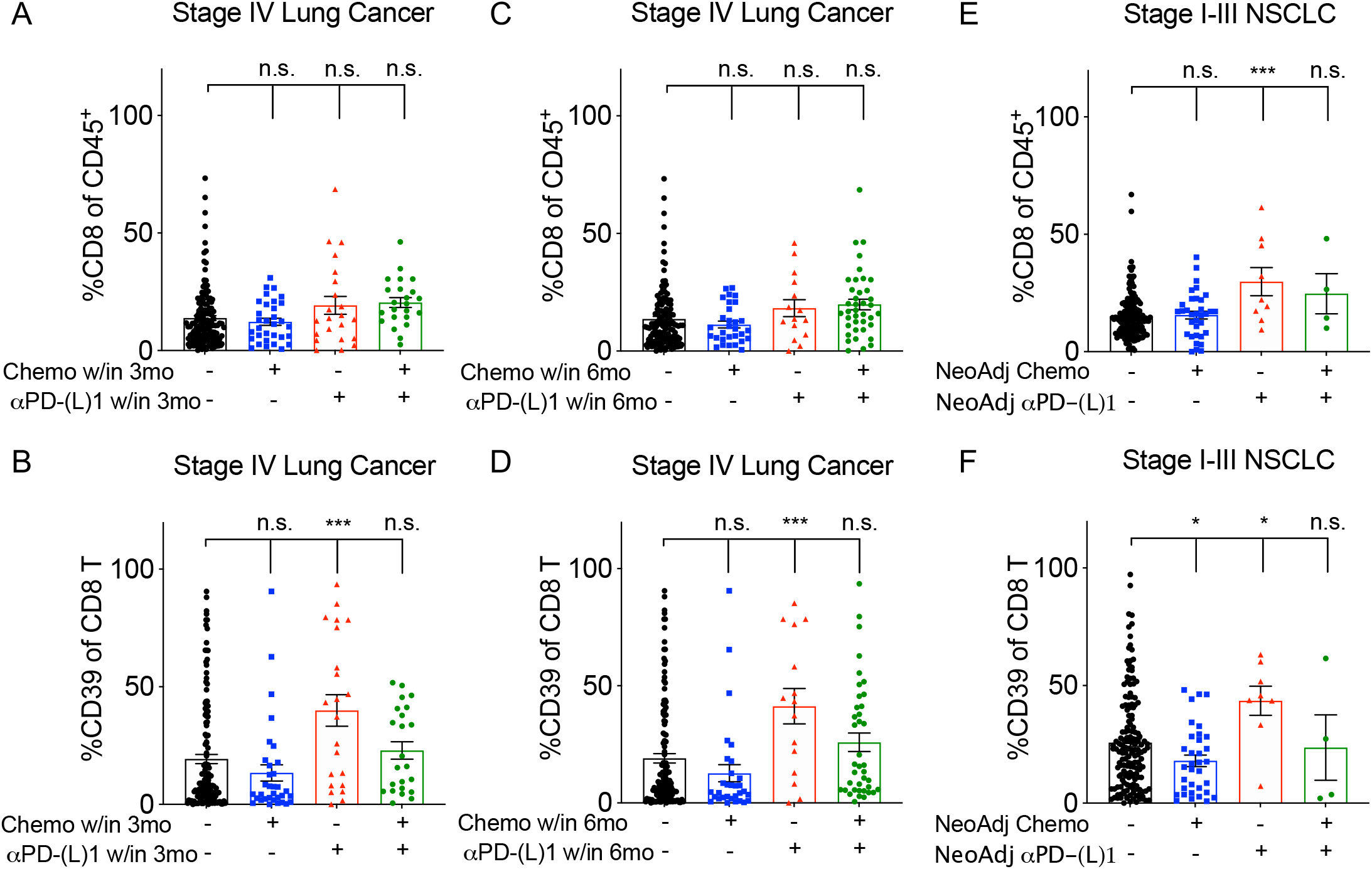
PD-1 axis blockade increases the frequency of CD39^+^ CD8 tumor-infiltrating lymphocytes. A-B) %CD8 (among CD45^+^) or %CD39 (among CD8 T cells) from 218 biospecimens obtained from patients with stage IV metastatic disease that did or did not receive chemotherapy or ICB therapy in the prior 3 months. Statistical significance was assessed by two-tailed student’s t-test. C-D) %CD8 (among CD45^+^) or %CD39 (among CD8 T cells) from 218 biospecimens obtained from patients with stage IV metastatic disease that did or did not receive chemotherapy or ICB therapy in the prior 6 months. Statistical significance assessed by two-tailed student’s t-test. E-F) %CD8 (among CD45^+^) or %CD39 (among CD8 T cells) from 208 biospecimens obtained from patients with early stage (stage I-III) NSCLC. Statistical significance was assessed by two-tailed student’s t-test. *p<0.05, ***p<0.001.

Since the stage IV lung cancer cohort is quite heterogeneous, we subsequently examined 208 early-stage (stage I-III) NSCLC biospecimens from patients that did or did not receive neoadjuvant therapy prior to sample collection. Among early-stage patients with NSCLC, we again did not observe differences in total CD8 infiltration with preceding neoadjuvant chemotherapy; however, neoadjuvant immunotherapy did increase total CD8 infiltration (**Fig. 3E**). CD8 T cells from patients who underwent neoadjuvant chemotherapy had reduced CD39 levels compared to patients without neoadjuvant therapy; in contrast, in patients who underwent neoadjuvant ICB, we observed an increase in CD39 expression on CD8 T cells (**Fig. 3F**). Thus, across two clinical cohorts, we did not find evidence for ‘immunogenic cell death’ with standard cytotoxic chemotherapy utilized for lung cancer; however, we did discover that ICB exposure was associated with an enhanced infiltration of CD39^+^ CD8 T cells. This is consistent with recent appreciation that ICB results in clonal expansion and infiltration of tumor-reactive CD8 T cells (*32*).

### CD39 and PD-1 expression on CD8 T cells portends benefit from ICB therapy

We next sought to evaluate whether CD39 on CD8 T cells had prognostic significance in our dataset. We first assessed whether CD39 levels at the time of resection in 188 patients with stage I-IIIA lung cancer resulted in differential recurrence-free survival (RFS). Since CD39 levels are modulated by prior ICB (**Fig 3F**), we excluded patients who had previously received neoadjuvant ICB in this analysis. Across this early-stage resection cohort, we did not observe differences in RFS based on total CD8 T cells or %CD39 among CD8 T cells utilizing mean cutoff values from the total 440 patient dataset (**Fig. 4A-C, Fig. S4A-C**). The data for CD8 T cells is in line with a recently published prospective cohort of early-stage NSCLC resection specimens which showed no differences in clinical outcomes based on total CD8 abundance (*33*). Similarly, neither total CD8 T cells or %CD39 among CD8 T cells resulted in differential progression-free survival (PFS) in the 26 stage IV lung cancer patients undergoing cytotoxic chemotherapy without ICB (**Fig. 4D, S4D**). Hence, CD39 expression on CD8 T cells was not prognostic for lung cancer in our dataset.

**Figure 4.**
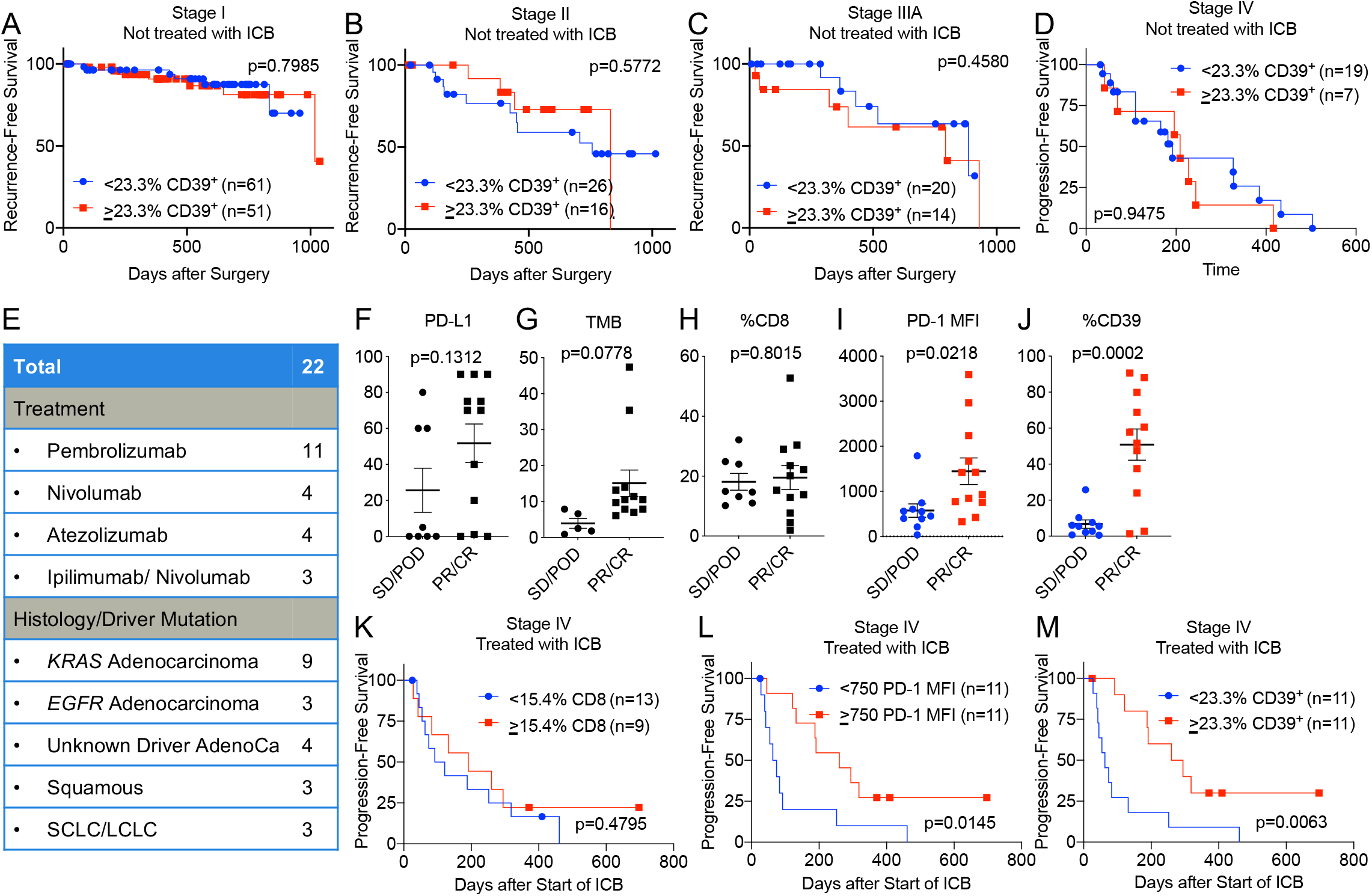
CD39 expression among CD8 tumor-infiltrating lymphocytes is not prognostic, but portends a beneficial outcome to immune checkpoint blockade in lung cancer. A-C) Kaplan-Meier curve of recurrence-free survival after resection with above or below mean %CD39 (among CD8 T cells) for stage I, stage II, and stage IIIA lung cancer patients who did not receive ICB. Statistical significance determined by Mantel-Cox test. D) Kaplan-Meier curve of progression-free survival after resection with above or below mean %CD39 (among CD8 T cells) for stage IV lung cancer patients who did not receive ICB. Statistical significance determined by Mantel-Cox test. E) Cohort of stage IV lung cancer patients who received ICB monotherapy. F-J) Tumor proportion score for PD-L1, tumor mutation burden, %CD8 (among CD45^+^), PD-1 mean fluorescence intensity (MFI) and % CD39 (among CD8 T cells) among non-responders (stable disease/progression of disease, SD/POD) or responders (partial response/complete response, PR/CR) to ICB in the cohort described in E). K-M) Kaplan-Meier survival curve of progression-free survival patients described in cohort E based on stratification for CD8, PD-1, and CD39. Statistical significance was determined by Mantel-Cox test.

We next asked whether we could find evidence that CD39 levels on CD8 T cells was associated with response to ICB. There were 22 patients in our cohort from whom we had obtained clinical biospecimens prior to or within three weeks of commencing ICB therapy (**Fig. 4E**). Although the tumoral PD-L1, TMB, and %CD8 among DAPI^-^ cells were not different between non-responders (stable disease/progression of disease, SD/POD) and responders (partial response/complete response, PR/CR) to ICB, we observed that responders had a higher level of PD-1 and CD39 on CD8 T cells (**Fig. 4F-J**). While total CD8 infiltration did not differentiate PFS in these patients (**Fig. 4K**), patients with higher levels of PD-1 or CD39 on CD8 T cells had an improved PFS (p=0.0145 for PD-1 and p=0.0063 for CD39, **Fig. 4L-M**). Thus, consistent with prior reports in lung cancer (*2, 34, 35*), levels of baseline exhausted CD8 T cells are associated with clinical benefit from ICB therapy.

## DISCUSSION

Lung cancer therapy is currently guided by tumor cell-intrinsic features, including histology, driver alterations, PD-L1 and TMB. Integration of tumor cell-extrinsic features such as the immune infiltrate can potentially lead to improved therapeutic options for patients with lung cancer. This was exemplified by the recent report of a ‘lung cancer activation module’, consisting of activated T cells, plasma cells, and macrophages, that served as a predictor of response to immunotherapy (*36*). Since CD8 T cells are the critical effectors in ICB therapy, features of CD8 T cells could also potentially serve as a non-redundant biomarker of response to ICB. Although methods to empirically verify tumor reactivity have improved substantially in sensitivity and throughput (e.g. MANIFEST (*37*) and 4-1BB assay (17)), there remains a need in certain contexts for simple and reliable methods to distinguish and quantify the tumor-reactive CD8 sub-population from bystanders without cognate TCR reactivity. We observed from single-cell profiling of heterogeneous lesions that CD8 T cells expressing high levels of CD39 are enriched for features of exhaustion, tumor reactivity and clonal expansion. Motivated by this finding and recent reports that empirically demonstrated that tumor-reactive CD8 T cells express CD39 (*2, 12, 13, 16, 17, 25, 38*), we profiled CD39 on 440 lung cancer biospecimens obtained at MSKCC. We observed that CD39 on CD8 T cells is expressed more highly in tobacco-associated lung cancer genotypes and poorly correlates with tumor TMB and PD-L1. Hence, CD39 is a biomarker that is non-redundant to tumoral features of lung cancer. CD39 is likely dependent on other variables not captured in our dataset, including T cell repertoire, HLA subtype, peptide binding, and likely the intersection of these three highly diverse attributes. CD39 is upregulated on CD8 T cells in the tumor microenvironment relative to the periphery and is generally stable across space and time. These data nominate CD39-guided strategies for TCR-based therapeutics against these targets (*39*). Finally, we demonstrate that although the level of CD39 on CD8 T cells was not prognostic in any of the patient cohorts that we examined, it was associated with clinical benefit from ICB therapy.

There have been conflicting data regarding whether the presence of CD8 T cells expressing markers of exhaustion can predict clinical benefit from ICB. Single-cell RNAseq of tumors from metastatic melanoma revealed that a CD8 T cell state called CD8_B, which is enriched in *ENTPD1* (CD39), *PDCD1* (PD-1), *CD38, HAVCR2* (TIM-3), and *LAG3*, is preferentially expressed in non-responders to ICB (9). Also, the use of multiplex immunofluorescence staining in lung cancer suggests that an ‘effector burnt out’ (EBO) CD8 T effector population preferentially expressing PD-1 and LAG-3 was enriched in patients who did not derive durable benefit from ICB and that these patients had reduced overall survival (*40*). Notably, CD39 was not a selective feature of the EBO CD8 T cells. On the other hand, lung cancer patients with high pre-treatment levels of PD-1 by flow cytometry or immunofluorescence demonstrated a greater response rate and PFS (*2, 34*). Moreover, high levels of intratumoral CD39^+^ CD8 T cells is associated with response to ICB therapy in lung cancer (*35*). These findings dovetail with other studies that revealed that high levels of CD8 T cells with features of exhaustion prior to ICB are associated with improved clinical benefit in colorectal cancer and ER^+^ breast cancer (*41, 42*). Our data confirm that the PD-1 and CD39 on CD8 T cells are both predictive of improved outcomes from ICB therapy.

Two recent reports highlight that the predictive significance of CD39 for immunotherapy is context dependent. Caushi et. al. demonstrated that although CD8 T cells that recognize neoantigen mutations express CD39, neoantigen-reactive CD8 T cells that expressed higher levels of CD39 were associated with inferior pathological responses (16). Krishna et. al. similarly showed that neoantigen-reactive CD8 T cells in *ex vivo* expanded TILs prior to infusion preferentially expressed CD39 and CD69, but improved clinical outcomes were observed in patients whose pre-infusion products harbored a high number of CD39^-^ CD69^-^ TILs (*43*). One explanation for this apparent paradox may be that the relative timing of the evaluation of exhaustion likely influences the predictive capability of this biomarker since our data demonstrates that CD39 increases after ICB. Notably, in the two cases highlighted above, the correlation of CD39 with worse outcomes was read out after immune based intervention (i.e. ICB or *ex vivo* expansion followed by adoptive cell transfer). It is plausible that pre-treatment/early on-treatment exhaustion is reflective of a mobilizable pool of tumor-reactive T cells that portends a favorable response to ICB; conversely, exhaustion after a course of ICB might be indicative of a persistent hypofunctional state that is associated with a worse clinical outcome.

There are several limitations of this study that we acknowledge. First, the data are derived from patients treated at a single center. Second, while we profiled CD39 on CD8 T cells, our data does not capture the likely contribution/modulation of other immune cell types such as B cells, innate lymphoid cells, and myeloid cells. Lastly, our cohort to assess the predictive significance of CD39 expression on CD8 T cells in stage IV lung cancer treated with ICB monotherapy is limited to 22 patients despite >3 years of fresh tissue collection. Part of this limited sample size is attributable to the routine incorporation of cytotoxic chemotherapy to ICB for frontline treatment of the majority of patients with lung cancer. Despite these limitations, the data presented here contribute substantially to our understanding of CD39 on CD8 T cells in the context of lung cancer, including factors that modulates its pattern of expression. Our study identifies CD39 on CD8 T cells as a biomarker that is non-redundant to TMB and PD-L1, and which can be readily captured on clinical samples to serve as a proxy of the tumor-reactive CD8 T cell population. Subsequent studies can also leverage CD39 to enrich for TCR candidates to evaluate for TCR-based immunotherapies.

## ACKNOWLEDGEMENTS

We are grateful for experimental support from the MSKCC Molecular Cytology Core Facility, Flow Cytometry Core Facility, and Integrated Genomics Operation Core (funded by the NCI Cancer Center Support Grant (CCSG, P30 CA08748), Cycle for Survival, and the Marie-Josée and Henry R. Kravis Center for Molecular Oncology). We are grateful for manuscript editing provided by Dr. Clare Wilhelm and Reeja Thomas.

## Funding

This research was funded in part through the NIH NCI CCSG P30 CA008748, NCI R01 CA056821, U24 CA213274, P01 CA129243, R01 CA197936, R37 CA259177 (CAK), Cancer Research Institute CRI3176 (CAK), Stony-Wold Herbert Fund, International Association of Lung Cancer Research (IASLC)/ International Lung Cancer Foundation (ILCF), the MSKCC Society Grant, the Ludwig Collaborative and Swim Across America Laboratory, the Emerald Foundation, the Parker Institute for Cancer Immunotherapy at MSKCC, the Department of Medicine at MSKCC, and Stand Up To Cancer (SU2C)-American Cancer Society Lung Cancer Dream Team Translational research grant (SU2C-AACR-DT17-15). AC was supported by an MSKCC Investigational Cancer Therapeutics Training Program fellowship (T32 CA-009207) and Clinical Investigator Award from National Cancer Institute (K08 CA-248723).

## Disclosure

CAK received research grant support from Kite/Gilead; is on the Scientific and/or Clinical Advisory Boards of Achilles Therapeutics, Aleta BioTherapeutics, Bellicum Pharmaceuticals, Catamaran Bio, Obsidian Therapeutics, and T-knife, and has performed consulting services for Bristol Myers Squibb, PACT Pharma, and Roche/Genentech. CAK is a co-inventor on patent applications related to TCRs targeting public neoantigens unrelated to the current work. MDH received research grant from BMS; personal fees from Achilles, Arcus, AstraZeneca, Blueprint, BMS, Genentech/Roche, Genzyme, Immunai, Instil Bio, Janssen, Merck, Mirati, Natera, Nektar, Pact Pharma, Regeneron, Shattuck Labs, Syndax, as well as equity options from Arcus, Factorial, Immunai, and Shattuck Labs. A patent filed by MSKCC related to the use of tumor mutational burden to predict response to immunotherapy (PCT/US2015/062208) is pending and licensed by PGDx. JDW is a consultant for Adaptive Biotech, Amgen, Apricity, Ascentage Pharma, Arsenal IO, Astellas, AstraZeneca, Bayer, Beigene, Boehringer Ingelheim, Bristol Myers Squibb, Celgene, Chugai, Daiichi Sankyo, Dragonfly, Eli Lilly, Elucida, F Star, Georgiamune, Idera, Imvaq, Kyowa Hakko Kirin, Linneaus, Maverick Therapeutics, Merck, Neon Therapeutics, Polynoma, Psioxus, Recepta, Takara Bio, Trieza, Truvax, Trishula, Sellas, Serametrix, Surface Oncology, Syndax, Syntalogic, and Werewolf Therapeutics. JDW has received grant/research support from Bristol Myers Squibb and Sephora. JDW has equity in Tizona Pharmaceuticals, Adaptive Biotechnologies, Imvaq, Beigene, Linneaus, Apricity, Arsenal IO, and Georgiamune. JDW is a co-inventor on patent applications related to heteroclitic cancer vaccines and recombinant poxviruses for cancer immunotherapy. JDW and TM are co-inventors on patent applications related to CD40 and in situ vaccination (PCT/US2016/045970). TM is a consultant for Immunos Therapeutics and Pfizer. TM is a cofounder of and equity holder in IMVAQ Therapeutics. TM receives research funding from Bristol-Myers Squibb, Surface Oncology, Kyn Therapeutics, Infinity Pharmaceuticals, Peregrine Pharmaceuticals, Adaptive Biotechnologies, Leap Therapeutics, and Aprea Therapeutics. TM is an inventor on patent applications related to work on oncolytic viral therapy, alpha virus–based vaccine, neoantigen modeling, CD40, GITR, OX40, PD-1, and CTLA-4. C.M.R. has consulted regarding oncology drug development with AbbVie, Amgen, Ascentage, AstraZeneca, BMS, Celgene, Daiichi Sankyo, Genentech/Roche, Ipsen, Loxo and PharmaMar and is on the scientific advisory boards of Elucida, Bridge and Harpoon.

## TABLE LEGENDS

**Table S1**. Clinical features of four lung cancer clinical biospecimens that underwent single cell CITE/TCR/RNA sequencing.

**Table S2**. Differentially expressed genes for CD8, CD4, T cell NOS, and Treg clusters from single cell CITE/TCR/RNA sequencing.

**Table S3**. Differentially expressed genes for CD39^hi^ CD8 T cells compared to all other CD8 T cells from single cell CITE/TCR/RNA sequencing.

**Table S4**. List of gene signatures used for scoring.

**Table S5**. Clinical and molecular features of 440 patient lung cancer cohort.

**Table S6**. Median and mean values for %CD39 on CD8 T cells among lung cancer subtypes.

**Table S7**. Linear regression multivariate analysis of %CD39 on CD8 T cells.

**Table S8**. Lasso regression multivariate analysis of %CD39 on CD8 T cells.

---

**Figure S1.**
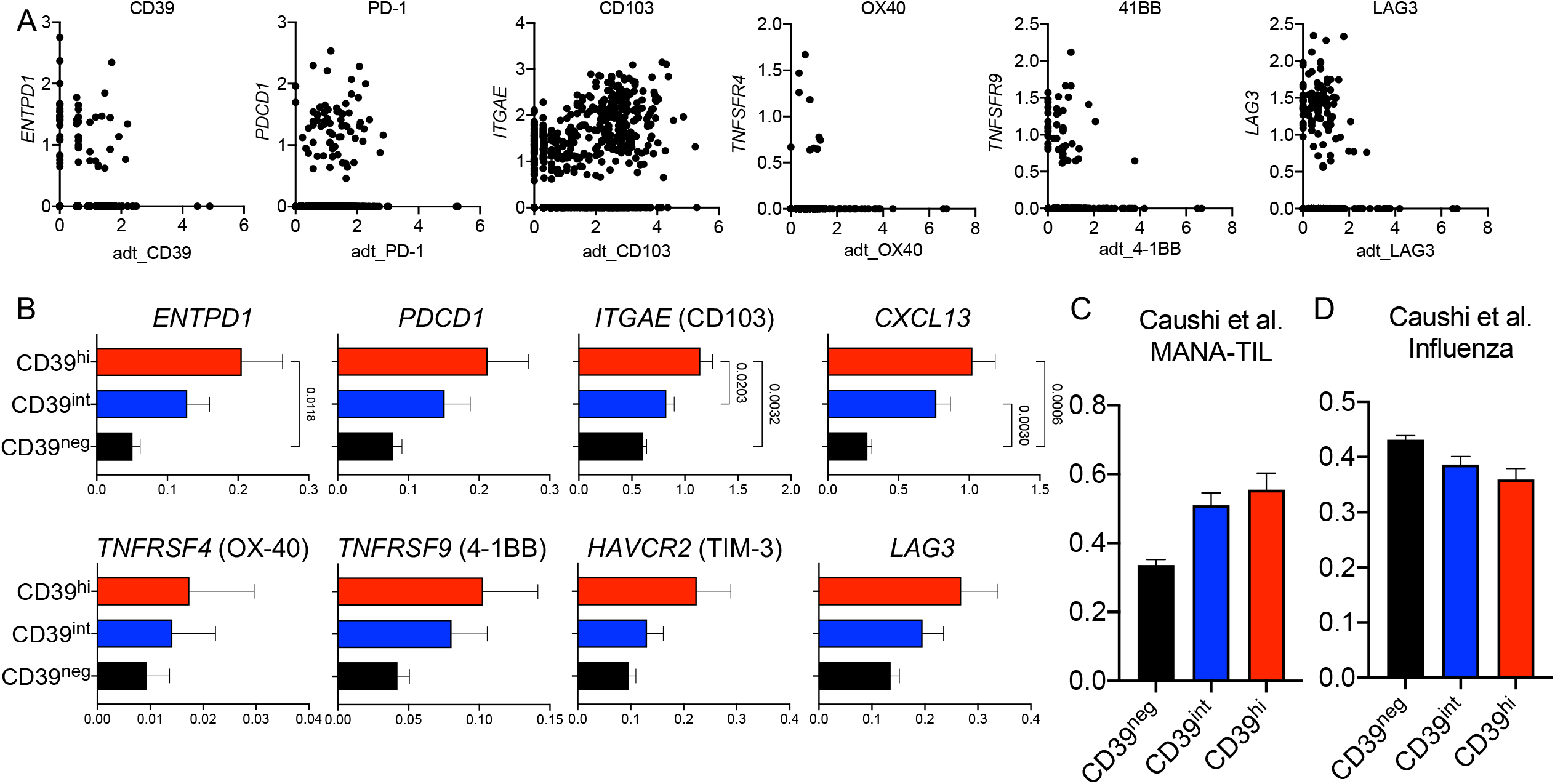
Single-cell CITE/TCR/RNA-sequencing reveals that CD39^hi^ CD8 T cells are enriched for features of tumor, but not viral, reactivity. A) Correlation of transcriptional expression (y axis) with protein expression (x axis) for indicated genes. B) Pooled transcript levels (scaled counts) of indicated genes in CD39^hi^, CD39^int^, and CD39^neg^ CD8 T cells as classified by adt_CD39 levels. C-D) Scaled scores for MANA-TIL and influenza-reactivity gene signatures (see Table S4 for full gene list). Statistical significance was determined by two-way ANOVA with Tukey’s multiple comparisons test and p value is indicated if < 0.05.

**Figure S2.**
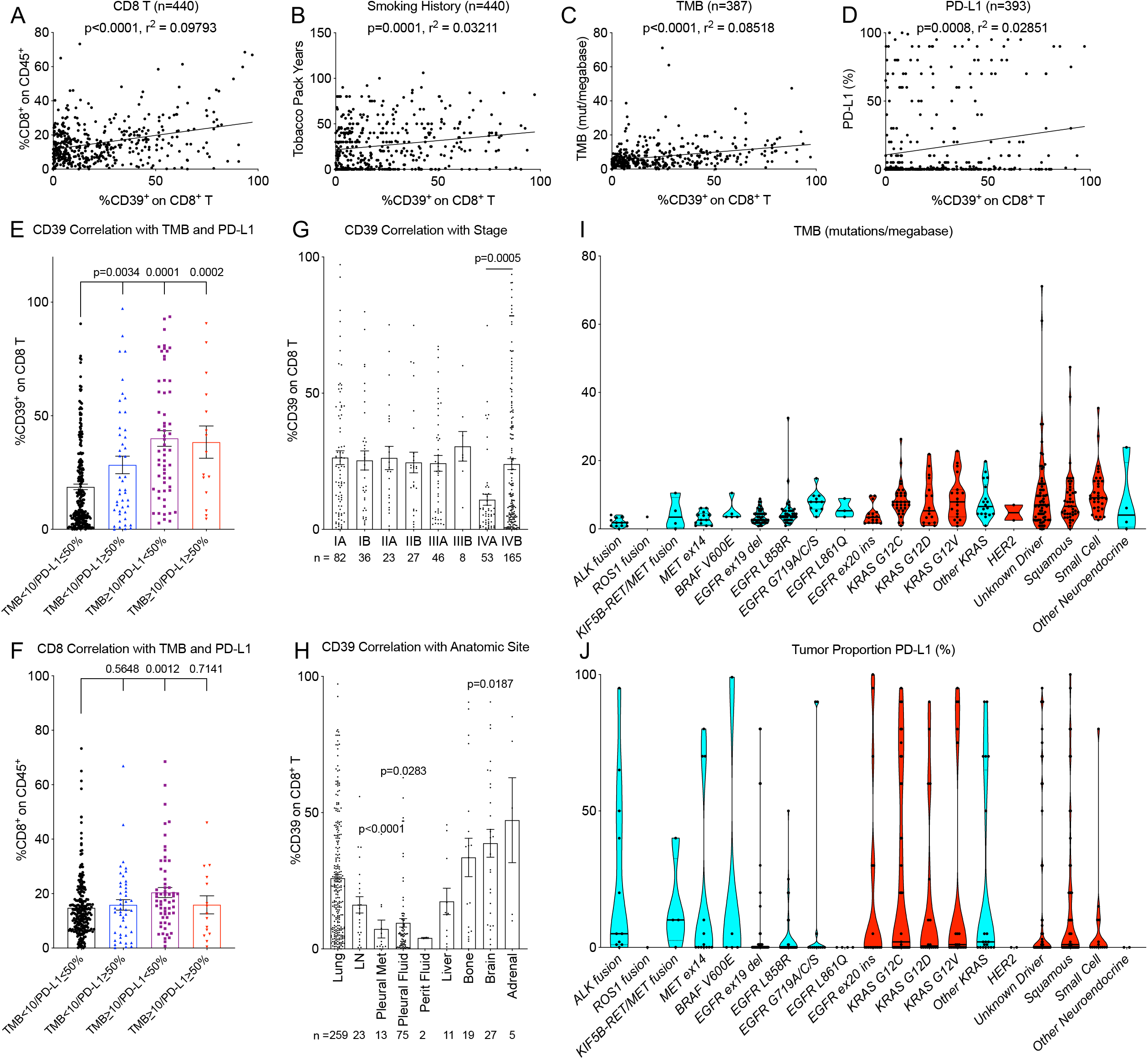
Clinical and molecular correlates of CD39 expression on CD8 tumor-infiltrating lymphocytes. A-D) Correlation of %CD39 on CD8 T cells with %CD8 (among CD45^+^), tumor mutation burden, PD-L1 tumor proportion score, and smoking history. E-F) %CD39^+^ (among CD8 T cells) and %CD8 (among CD45^+^) on four subgroups based on cutoffs for TMB and PDL-1 in the 440-sample cohort. Statistical significance was assessed by two-tailed student’s t-test. G) Correlation of %CD39 on CD8 T by lung cancer stage. H) Correlation of %CD39 on CD8 T by anatomic site of biospecimen. I-J) Correlation of tumor mutation burden and PD-L1 tumor proportion score with various histological subtypes/driver mutation categories among 440-sample cohort. Red bars indicate histological subtypes/driver mutations with above average %CD39 values.

**Figure S3.**
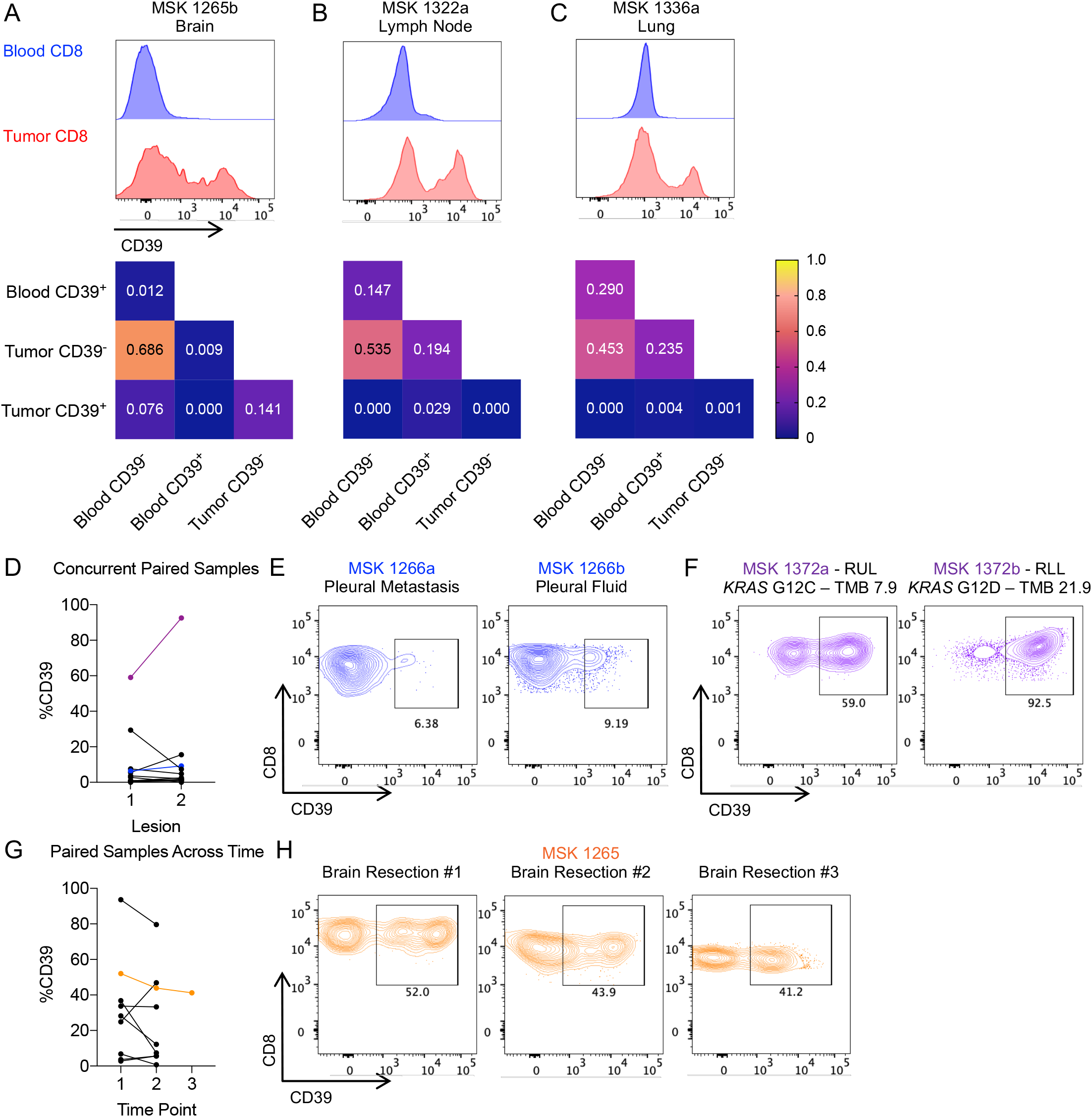
CD39 is upregulated in the tumor microenvironment and is consistent across space and time. A-C) Top row depicts flow cytometry histogram plots of CD39 levels on DAPI^-^ CD45^+^ CD3^+^ CD8^+^ T cells from peripheral blood or matched tumor lesion from three patients with lung cancer. Bottom row depicts Morisita indices depicting TCR-β overlap among sorted CD39^-^ and CD39^+^ fractions of blood and tumor CD8 T cells. D) %CD39 among CD8 T cells from two spatially distinct lesions that were sampled concurrently in the same patient (n=12). Blue line indicates MSK 1266 whose CD39 expression is depicted in E. Purple line indicates MSK 1372 depicted in F. E) Flow cytometry plots of CD39 expression on CD8 T cells on two concurrent pleural lesions in MSK 1266. F) Flow cytometry plots of CD39 expression on CD8 T cells in two concurrent spatially distinct lung lesions in MSK 1372. G) %CD39 among CD8 T cells from two lesions from the same anatomic site that were sampled at temporally distinct times in the same patient (n=9). Orange line indicates MSK 1265 whose CD39 expression is depicted in H. H) Flow cytometry plots of CD39 expression on CD8 T cells over the course of three brain resection lesions in MSK 1265.

**Figure S4.**
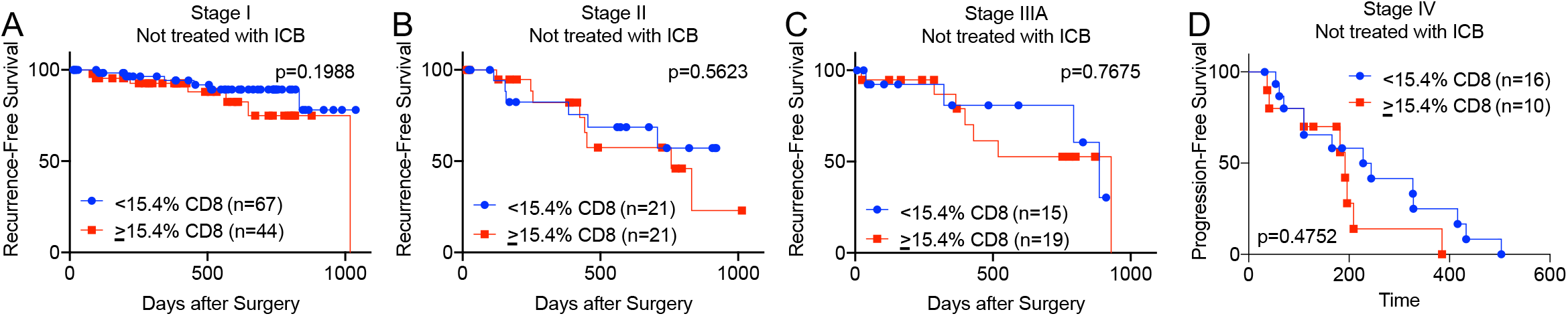
Total CD8 tumor-infiltrating lymphocytes is not prognostic for survival in lung cancer. A-C) Kaplan-Meier curve of recurrence-free survival stratified by %CD8 (among CD45^+^) for stage I, stage II, and stage IIIA lung cancer patients who did not receive ICB. Statistical significance was determined by Mantel-Cox test. D) Kaplan-Meier curve of progression-free survival stratified by %CD8 (among CD45^+^) for stage IV lung cancer patients who did not receive ICB. Statistical significance was determined by Mantel-Cox test.

## Notes

### Competing Interest Statement

Please check manuscript file.

### Summary of Updates

V3

